# Decoding spoken English phonemes from intracortical electrode arrays in dorsal precentral gyrus

**DOI:** 10.1101/2020.06.30.180935

**Authors:** Guy H. Wilson, Sergey D. Stavisky, Francis R. Willett, Donald T. Avansino, Jessica N. Kelemen, Leigh R. Hochberg, Jaimie M. Henderson, Shaul Druckmann, Krishna V. Shenoy

## Abstract

**Objective:** To evaluate the potential of intracortical electrode array signals for brain-computer interfaces (BCIs) to restore lost speech, we measured the performance of classifiers trained to discriminate a comprehensive basis set for speech: 39 English phonemes. We classified neural correlates of spoken-out-loud words in the “hand knob” area of precentral gyrus, which we view as a step towards the eventual goal of decoding attempted speech from ventral speech areas in patients who are unable to speak.

**Approach:** Neural and audio data were recorded while two BrainGate2 pilot clinical trial participants, each with two chronically-implanted 96-electrode arrays, spoke 420 different words that broadly sampled English phonemes. Phoneme onsets were identified from audio recordings, and their identities were then classified from neural features consisting of each electrode’s binned action potential counts or high-frequency local field potential power. We also examined two potential confounds specific to decoding overt speech: acoustic contamination of neural signals and systematic differences in labeling different phonemes’ onset times.

**Main results:** A linear decoder achieved up to 29.3% classification accuracy (chance = 6%) across 39 phonemes, while a recurrent neural network classifier achieved 33.9% accuracy. Parameter sweeps indicated that performance did not saturate when adding more electrodes or more training data, and that accuracy improved when utilizing time-varying structure in the data. Microphonic contamination and phoneme onset differences modestly increased decoding accuracy, but could be mitigated by acoustic artifact subtraction and using a neural speech onset marker, respectively.

**Significance:** The ability to decode a comprehensive set of phonemes using intracortical electrode array signals from a nontraditional speech area suggests that placing electrode arrays in ventral speech areas is a promising direction for speech BCIs.

## 1. Introduction

Neurological disorders such as stroke and amyotrophic lateral sclerosis (ALS) take away the ability to speak from millions worldwide (LaPointe and Stierwalt, 2018). In the case of ALS, loss of speech is often considered the worst aspect of disease progression, necessitating augmentative communication within 1-2 years of symptom onset (Makkonen et al., 2018). Noninvasive assistive technologies such as sip-and-puff interfaces and eye tracking have helped re-establish communication, but suffer from low information transfer rates and limited expressivity (Fager et al., 2019; Tai et al., 2008). In contrast, speech is amongst the most intuitive and fastest means of communication, with conversational speech conveying roughly 150 words per minute (Chang and Anumanchipalli, 2020). Brain-computer interfaces (BCIs) are an emerging technology through which a person’s movement intentions are read out from their neural signals (Slutzky, 2019; Tam et al., 2019). A high-performance speech BCI that could accurately identify what people with speech loss want to say would therefore substantially improve patients’ quality of life.

To gain direct access to the relevant neural signals, ongoing efforts to build speech BCIs (reviewed in (Chakrabarti et al., 2015; Herff and Schultz, 2016; Martin et al., 2018; Rabbani et al., 2019) have often used invasive electrocorticography (ECoG) measurements, in which many electrode disks distributed across the cortical surface sample the aggregative activity of many tens of thousands of neurons under each recording site. This approach has been used in discrete classification of small building blocks such as phonemes, syllables, words, and sound snippets (Bouchard and Chang, 2014; Herff et al., 2015, 2019; Kellis et al., 2010; Martin et al., 2014; Mugler et al., 2014; Ramsey et al., 2018; Salari et al., 2018). Recent work has also used neural networks to learn nonlinear mappings that transform neural signals into a variety of outputs such as acoustic speech features (Akbari et al., 2019; Angrick et al., 2019), including by way of intermediate articulatory kinematics (Anumanchipalli et al., 2019), discrete targets such as phonemes or syllables (Livezey et al., 2019), and entire phrases and sentences (Makin et al., 2020; Moses et al., 2019).

While the most advanced speech BCIs to date are ECoG-based, other recording modalities are also being investigated. These include non-surgical methods such as electroencephalography (Nguyen et al., 2018) and magnetoencephalography (Dash et al., 2020), stereotactic electroencephalography to access field potentials from deeper brain structures (Herff et al., 2020), and intracortical electrodes capable of recording neuronal spiking activity (Brumberg et al., 2011; Guenther et al., 2009). The intracortical approach, which can in principle sample from many distinct sources of information (individual neurons or small groups of neurons), is particularly promising for providing the sufficiently high neural bandwidth necessary to restore conversational speech. Intracortical BCIs that use multielectrode array signals to decode attempted arm and hand movements have demonstrated the highest performance to date for controlling computer cursors (Pandarinath et al., 2017), high degree-of-freedom robotic arms (Collinger et al., 2013), and the user’s own arm muscles (Ajiboye et al., 2017), with no prior user training (Brandman et al., 2018). In contrast, intracortical BCIs for decoding speech are far less mature. Several studies have examined speech production using small numbers of intracortical recording sites in lateral temporal lobe (Creutzfeldt et al., 1989; Tankus et al., 2012), ventral motor cortex (Brumberg et al., 2011; Guenther et al., 2009), orbitofrontal cortex (Tankus et al., 2012), and subthalamic nucleus (Lipski et al., 2018; Tankus and Fried, 2019). Ninety-six-electrode Utah arrays, of the type used in the aforementioned arm and hand BCIs, have been placed into superior temporal gyrus to decode heard speech in monkeys (Heelan et al., 2019) and both heard and spoken speech in a person (Chan et al., 2014). However, these previous studies did not use large numbers of electrodes to record from motor areas of cortex, which is the approach that has proved promising for restoring arm and hand function.

We are positioned to start to address this gap after having recently found speech-related activity in the dorsal “hand/knob” area of precentral gyrus (Stavisky et al., 2019; Willett et al., 2020a), where a number of groups already place Utah arrays.These signals could be used to discriminate between a small number of syllables and words with high accuracy (Stavisky et al., 2018, 2019, 2020). In the present study, we build on this discovery to further evaluate the potential of using intracortical signals from two Utah arrays to decode amongst 39 phonemes, which could be used as a comprehensive basis set in a speech BCI. Importantly, we recorded from the dorsal precentral gyrus of two people with tetraplegia who already had arrays placed as part of their participation in the BrainGate2 clinical trial, which has the primary goal of evaluating the safety of these intracortical BCI devices. We recognize and wish to highlight that this cortical area, which has a well-established role in volitional arm and hand movements (Ajiboye et al., 2017; Bouton et al., 2016; Collinger et al., 2013; Downey et al., 2018a; Hochberg et al., 2006, 2012; Pandarinath et al., 2017; Rastogi et al., 2020; Wodlinger et al., 2015) and to a lesser extent movements of other body parts (Stavisky et al., 2019; Willett et al., 2020a), is likely sub-optimal for decoding speech. Nonetheless, this research context also presents a rare opportunity to evaluate the feasibility of decoding speech production from many simultaneously recorded intracortical signals, using ground truth data from people who can still speak, without additional risk. Our guiding motivation is that the results of this investigation should be viewed as a lower bound on the intracortical performance that ought to be possible, and that higher performance will be possible with arrays specifically placed in ventral speech cortex where previous ECoG studies have shown strong speech production-related modulation (Bouchard et al., 2013; Chartier et al., 2018; Herff et al., 2019; Kellis et al., 2010; Mugler et al., 2018; Takai et al., 2010).

We recorded neural signals as the participants spoke out loud a set of 420 different short words. This set included most of the same spoken words as a previous ECoG study (Mugler et al., 2014), which allowed us to make a well-matched comparison between decoding Utah array signals and this previous ECoG result. We first examined whether neural firing rates changed when the participant spoke just one or a handful of phonemes (“sparse tuning”), or across speaking many phonemes (“broad tuning”). This measurement is important for assessing the viability of intracortical speech BCIs, which in the near-term are likely to be restricted to hundreds (not thousands) of electrodes located within a relatively localized area of cortex. We found that activity recorded on these electrodes had broad tuning; this encoding makes it more likely that signals related to producing all the phonemes will be observable from a limited number of electrodes, and also has previously been shown to be more robust and decodable (Abbott and Dayan, 1999). Next, we trained both conventional and deep learning classifiers to predict which of 39 phonemes was being spoken, using the available neural signals. We found that decoding performance was better using high-frequency local field potentials (HLFP) rather than threshold crossing spikes (i.e., action potentials from one or potentially several neurons near an electrode tip detected when the measured voltage drops below a set threshold), and was competitive with previous ECoG decoding performance (Mugler et al., 2014). Encouragingly, subsampling the data used for decoder training indicated that speech BCI performance is likely to improve as more training data and electrode recording sites become available.

While there are many advantages to using overt speaking data to establish proof-of-feasibility for a speech BCI, this widely used model (Angrick et al., 2019; Anumanchipalli et al., 2019; Bouchard and Chang, 2014; Herff et al., 2015, 2019; Kellis et al., 2010; Livezey et al., 2019; Makin et al., 2020; Moses et al., 2019; Mugler et al., 2014; Pailla et al., 2016; Ramsey et al., 2018; Salari et al., 2018; Suppes et al., 1997) also introduces potential confounds. Here we investigated two potential limitations which have received little prior attention. The first is that using the recorded audio signal to detect voice onset may introduce systematic onset time biases across phonemes due to differences between when voice sounds become detectable versus when the speech articulators are being moved. This in turn can exaggerate neural differences between phonemes by artificially shifting what are actually condition-invariant neural signal components. The second confound, which was recently raised by Roussel and colleagues (Roussel et al., 2019), is that mechanical vibrations due to speaking might cause microphonic artifacts in the neural recordings. Our analyses suggest that while these confounds most likely do inflate speech decoding performance, their effects are not large. Furthermore, we introduce analysis methods that can be applied to neural data collected during overt speech to mitigate these confounds.

## 2. Methods

### 2.1 Participants

Research sessions were conducted with volunteer participants enrolled in the BrainGate2 pilot clinical trial (ClinicalTrials.gov Identifier: NCT00912041). The trial is approved by the U.S. Food and Drug Administration under an Investigational Device Exemption (Caution: Investigational device. Limited by Federal law to investigational use) and the Institutional Review Boards of Stanford University Medical Center (protocol #20804), Brown University (#0809992560), and Partners HealthCare / Massachusetts General Hospital (#2011P001036). The BrainGate2 trial’s purpose is to collect preliminary safety information and demonstrate feasibility that an intracortical BCI can be used by people with tetraplegia for communication and control of external devices; the present manuscript results from analysis and decoding of neural activity recorded during the participants’ engagement in research that is enabled by the clinical trial but does not report clinical trial outcomes.

Participant T5 is a right-handed man who was 65 years old at the time of the study. He was diagnosed with C4 AIS-C spinal cord injury eleven years prior to this study. T5 is able to speak and move his head, and has residual movement of his left bicep as well as trace movement in most muscle groups. Participant T11 is an ambidextrous man who was 35 years old at the time of the study. He was diagnosed with C4 AIS-B spinal cord injury eleven years prior to this study. T11 is able to speak and move his head. He has some residual movement in both arms. Both participants gave informed consent for this research and associated publications.

### 2.2 Many Words Task

The participants performed a simple visually prompted speaking task. On each trial they spoke one of 420 unique words that widely sample American English phonemes. The words used are an expanded set of the Modified Rhyme Test (House et al., 1963; Mines et al., 1978) and were previously used in several other studies (Angrick et al., 2019; Herff et al., 2019; Mugler et al., 2018). They include most, but not all, of the words used in Mugler and colleagues’ study (Mugler et al., 2014). These words were visually prompted (and subsequently spoken) one at a time. The participant was seated while fixating on a colored square centered on a computer screen in front of him. A trial started with an instruction period lasting 1.2 to 1.8 seconds, in which the central square was red, and white text above it instructed the word (e.g., ‘Prepare: “Word’, see **Fig. 1B**). The participant subsequently spoke the prompted word out loud following a go cue, which consisted of the square turning green, the text changing to “Go,” and an audible beep. The next trial began ~2.5 seconds later. Trials were presented in continuous blocks separated by short breaks, with the entire 420 word corpus divided across four blocks. Participant T5 performed three repetitions of each word (12 total blocks). Participant T11 performed 11 total blocks resulting in two to three repetitions of each word. Data from participant T5 were previously analyzed in (Stavisky et al., 2019), which examined the phonemic structure of trial-averaged, phoneme-aligned firing rates. The T11 data have not previously been reported.

**Figure 1.**
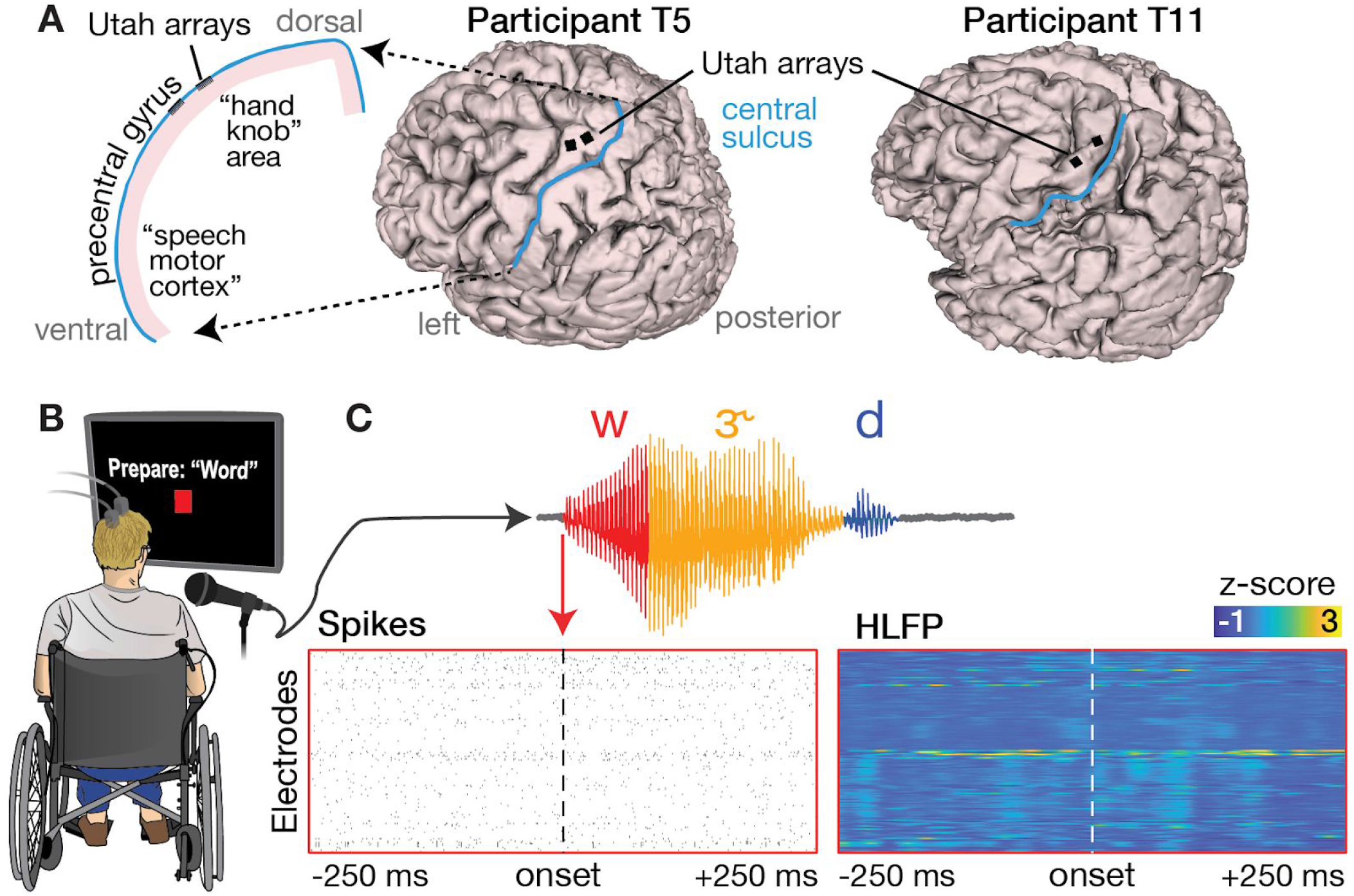
Neural data recorded during a word speaking task. (**A**) Array placements on 3D reconstructions of each participant’s brain. The left side illustration highlights that we recorded neural correlates of overt speech in a dorsal cortical area that is distinct from the ventral areas where speech production is typically decoded. (**B**) Illustration of the visually-prompted word speaking task. (**C**) Example phoneme segmentation of a word from the recorded audio. Below we show threshold crossing spikes and high frequency LFP (HLFP) for a 500 ms window rates centered on voice onset for this utterance of /w/.

Phoneme classes with at least 30 repetitions in the T5 dataset were included in the phoneme-specific analyses (e.g., classification), resulting in a set of 39 different phonemes. One phoneme in the T11 dataset (/ɔɪ/) had fewer repetitions (26) but was nonetheless included for consistency across the two participants. Two phonemes which otherwise would have had too few repetitions for inclusion were consolidated with a very similar phoneme with more repetitions: /ɔ/ (14 T5 utterances, 14 T11 utterances) was grouped with /ʌ/ (115 T5 utterances, 96 T11 utterances), and /ɚ/ (18 T5 utterances, 18 T11 utterances) was grouped with /ɝ/ (30 T5 utterances, 29 T11 utterances).

In the phoneme classification analyses shown in **Supplementary Figure 2**, we sought to match, as best we could, a set of words for comparing phoneme decoding with that of a previous ECoG phoneme decoding study by Mugler and colleagues (Mugler et al. 2014). Our word set already had 256 of the 320 words of that study; these we included in our ‘matched’ data subset. To fill out the remaining 64 words, we picked similar words from the 190 additional words in our overall dataset. Our substituted words have the same total number of phonemes as the missing words in the Mugler and colleagues report (Mugler et al., 2014), and typically differed in only one or two phonemes. Examples of these substitutions are “bill” (bɪl) → “bell” (bɛl), “late” (leɪt) → “loot” (lut), “mass” (mæs) → “mouse” (maʊs), “ray” (reɪ) → “reach” (ritʃ), “tip” (tɪp) → “type” (taɪp).

### 2.3 Intracortical electrode array recording

Participants T5 and T11 had two 96-electrode Utah arrays (Blackrock Microsystems Inc., Salt Lake City, USA) neurosurgically placed in the ‘hand knob’ area of their left dorsal motor cortex 29 months (T5) and 3 months (T11) prior to this study (**Fig. 1A**).

Neural signals were acquired with a NeuroPort™ system (Blackrock Microsystems Inc.), analog filtered from 0.3 Hz to 7,500 Hz, and digitized at 30,000 Hz at 16 bits/sample.

### 2.4 Neural signals

We common average referenced each electrode by subtracting the mean voltage across all other electrodes from its time series at each timepoint. Action potentials (spikes) were detected based on when voltage values became more negative than a threshold set at −3.5 × root mean square (RMS) of that electrode’s voltage. We did not spike sort these threshold crossing spikes (‘TCs’), which may capture action potentials from one or more neurons near the electrode tip, since here we were interested in decoding the neural state rather than characterizing the properties of putative single neurons (Ajiboye et al., 2017; Brandman et al., 2018; Collinger et al., 2013; Even-Chen et al., 2018; Oby et al., 2016; Pandarinath et al., 2017; Trautmann et al., 2019). The voltage time series were also bandpass-filtered using a 3rd order bandpass Butterworth causal filter from 125 to 5000 Hz to extract high-frequency local field potential (HLFP) signals, which have been previously shown to contain useful motor-related information (Pandarinath et al., 2017; Stavisky et al., 2018, 2019).

### 2.5 Audio recording

Audio recordings were made using a microphone (Shure SM-58), pre-amplified by ~60 dB (ART Tube MP Studio microphone pre-amplifier), and digitized at 30,000 Hz by the NeuroPort system via an analog input port.

Phoneme identities and onset times were manually labeled using the Praat software package (Boersma, 2001). The spoken words contained between two and six phonemes with a majority being three phonemes long (**Supplementary Fig. 1A**). **Supplementary Figure 1B,C** shows the distribution of phoneme audio durations. Although the same word prompt set was used for both participants, participant T5 completed one additional block of the many words task, and both participants occasionally misspoke or chose to speak different words that are spelled the same way (e.g., “tear”). Thus, the exact utterance count (T5: 3,840, T11: 3,467) and distribution of spoken phonemes differed between participants.

### 2.6 Neural feature extraction and classification models

HLFP activity was clipped at 100 x median activity for each electrode to lessen the impact of rare, large-amplitude electrical noise artifacts. HFLP and spike rasters were temporally smoothed using a moving average filter (20 milliseconds with 1 millisecond spacing), resulting in time series of binned TCs firing rates and HLFP power. Electrodes were then visually inspected and those with baseline firing rate nonstationarities or visible noise were excluded from the neural analyses; this resulted in 158 and 179 included electrodes for T5 and T11, respectively.

We trained (1) a multi-class logistic regression model and (2) a recurrent neural network (RNN) to predict phoneme identity from neural features using the Scikit-learn and Tensorflow (version 1.15.0) libraries respectively (Abadi et al., 2016; Pedregosa et al., 2011). Both models were trained in a cross-validated classification setup using multiple bins of activity per electrode as described below.

Except where otherwise mentioned, we extracted a 500 millisecond window of neural activity centered around the onset of each phoneme. For the logistic regression model, we averaged activity in this window within non-overlapping 50 millisecond bins (Mugler et al., 2014) and z-score normalized their magnitudes to account for different electrodes’ activity ranges. We then compressed the resulting feature set using principal components analysis (PCA, 75% variance retained). Z-score normalization mean and standard deviation (s.d.), and PCA coefficients were calculated from within each training fold and then applied to the test data.

For the RNN decoder analysis (see Section 2.8: *Recurrent neural network decoder*), we did not discard the 34 T5 electrodes as described above and instead used all 192 electrodes. Our reasoning was that even non-stationary or noisy electrodes could potentially be used by the RNN (Sussillo et al., 2016). In practice, however, this made little difference: the RNN decoder with the same electrode exclusions as the linear decoder performed at 32.4% accuracy, compared to 33.9% without excluding any electrodes.

### 2.7 Logistic regression for phoneme classification

The logistic regression model was trained using the Scikit-learn library (Pedregosa et al., 2011) in a leave-one-out cross-validation setup using default parameters except for the “lbfgs” optimization setting and L2 regularization value (λ = 100) to suppress overfitting. For multi-class classification, we used a one-vs-rest scheme where a different model is trained for each class to discriminate it against all others. For a given phoneme, model predictions are pooled and the highest positive class likelihood across models is taken as the decoder estimate.

We used 20-fold cross-validation for generating the phoneme classifications shown in **Figs. 4**, **5E**, and **5F**. We used leave-one-out cross-validation for the phoneme classifications shown in **Figs. 3A**, **3F**, **5D** and **Supplementary Fig. 2**. For all hyperparameter sweeps (**Fig. 3**), we used 10 repetitions of a 10-fold cross-validation procedure. Training data duration sweeps were performed by taking fractions of the overall phoneme data and then, for each such fraction, calculating the mean training data audio duration across training folds. This approach ensured that audio length estimates were not inflated by overlapping phoneme windows in the same training fold (e.g., if two consecutive phonemes are included), which would be the case if we had simply multiplied the number of phonemes used by our 500 millisecond window. The “number of electrodes” hyperparameter sweep was performed by randomly selecting a specific number of electrodes for both training and testing, and repeating this sampling 10 times for each number of electrodes.

To test for structure in the decoder’s misclassifications of T5’s neural data, we sorted our confusion matrix, where entry (i, j) corresponds to the percentage of times the *i*th phoneme in our decoding set is classified as the *j*th, by place of articulation (as in (Moses et al., 2019; Stavisky et al., 2019) and measured the difference between within- and between-group confusions. We then built a null distribution of expected differences if there were no underlying structure correlated with place of articulation in these errors by generating random partitions of our phonemes (keeping the same group sizes) and re-measuring the test statistic 10,000 times from these permuted data.

### 2.8 Recurrent neural network decoder

A recurrent neural network (RNN) was trained to predict phoneme identities from 1000 ms neural data snippets from each electrode, each subdivided into fifty 20 ms bins. The RNN was built as a single layer of 512 bidirectional gated recurrent units (GRUs) (Cho et al., 2014) implemented with the cuDNN library (Nvidia Corp., Santa Clara, USA). When training, the input data at each 20 ms time step was the 192-dimensional HLFP feature vector, while the supervised target output was a 39-dimensional ones-hot vector where the element corresponding to the phoneme associated with this data snippet was set to 1, and all other phonemes were set to 0 (**Supplementary Fig. 3A**). The training cost function also included a L2 regularization term to penalize large network weights (λ = 1e-5). During testing, for each phoneme utterance a new 192×50 neural data matrix was input to the RNN, and a 39×50 output matrix was read out, which consisted of the RNN’s predicted likelihoods (logits) that the input data came from each of the 39 possible phonemes, for each bin. The most likely phoneme during the last bin was chosen as the RNN’s final prediction for that snippet (**Supplementary Fig. 3B**). The RNN was trained and evaluated across 10 folds of the data, such that each spoken phoneme utterance appeared in the test set once.

Given the relative paucity of available Many Words Task speaking data compared to the data corpus sizes in typical machine learning applications, two additional training data augmentation methods were used to regularize the RNN and prevent it from overfitting on the training data. These were adapted from our group’s recent work using RNNs to decode the neural correlates of handwriting (Willett et al., 2020b) and conform to lessons learned from a previous iBCI RNN decoding study (Sussillo et al., 2016). First, white noise (s.d. = 0.1) was added to each time bin’s neural input feature vectors during training. Second, more slowly-varying artificial neural input feature drifts were added across the time bins of each snippet during training to make the RNN robust to potential nonstationarities in the neural data (Downey et al., 2018b; Jarosiewicz et al., 2015; Perge et al., 2013). These drifts had two components: a constant offset vector applied separately to each utterance (s.d. = 0.6), and a random walk across bins (s.d. = 0.02)

The RNN was trained using the “Adam” gradient descent optimization (Kingma and Ba, 2017) with a mini-batch size of 128 utterances, with no burn in steps, for 20,000 mini-batches. The learning rate decreased linearly from 0.01 (first mini-batch) to 0 (last mini-batch).

### 2.9 Demixed Principal Components Analysis (dPCA)

We used dPCA (Kobak et al., 2016) to decompose population activity into components reflecting variance that can be attributed to phoneme classes and time (“phoneme-dependent”) or time only (phoneme-independent). Specifically, we took a 1500 ms window centered on voice onset and decomposed it into low-rank approximations that captured phoneme class-dependent and independent variance. Spike rasters were convolved with a symmetrical 25 ms s.d. Gaussian and then averaged within 10 ms, non-overlapping bins before applying dPCA.

To ensure that dPCs were not fitting to noise, we adopted the cross-validation procedure described in (Willett et al., 2020a) to obtain 95% confidence intervals. We split our trials into ten separate folds to cross-validate dPC decompositions, thus avoiding overfitting to dataset noise. For each fold, all other folds are used to identify a dPCA decomposition. We then projected the held-out trials onto the identified dPCs. Geometrically, dPCA finds a linear subspace of a marginalized feature space where the axes correspond to a reconstruction error-minimizing subspace. Flipping the orientation of these vectors will preserve this subspace and hence also satisfy the optimization, but can confuse visualizations and comparisons across dPCA application instances. To avoid this issue and facilitate dPC comparisons across folds, we multiplied our components by −1 when this would result in a smaller angle between the *k*th principal components in a given marginalization.

We used the largest phoneme-independent neural component, which we refer to as the 1st component of the condition-invariant signal (CIS_1_) as in (Kaufman et al., 2016), to determine when phoneme production started from the neural data. This is in contrast to using acoustic data (**Fig. 4**). To do so, we trial averaged the neural activity for a given phoneme within each cross-validation fold and projected the resulting electrodes by time matrix onto the CIS_1_ neural dimension. This yields 10 CIS peak estimates (for 10 folds). The average CIS_1_ time course across these ten folds was used to determine each phoneme’s CIS_1_ peak (the time of maximum CIS_1_ value), which determined that phoneme’s neurally-derived onset.

### 2.10 Quantifying acoustic artifact and Linear Regression Reference (LRR) decontamination

To compare spectral content between recorded audio and electrodes (**Fig. 5A,B)**, we convolved the voltage time series of each electrode and also the audio channel with a 200 ms Hamming window and then computed the power spectral density (PSD) in non-overlapping bins using a short-time Fourier transform (as in Roussel *et al.* 2019). We isolated ‘voicing epochs’ in which to compare audio and neural power time series by sub-selecting time points with summed audio power (across all frequencies) in the top ~10% of values across all audio data. For spectrogram visualizations in **Fig. 5A**, sliding time windows overlap by 90%.

The LRR procedure (Young et al., 2018) to remove putative acoustic contamination from each electrodes’ signal began with fitting linear regression models that predicted each electrode’s voltage (bandpass-filtered in the 125-5000 Hz range using a third-order Butterworth filter) at a given time sample from every other electrode’s voltage at that time sample. Regression weights were fit separately within each block in order to increase reliability if there were across-block changes in the acoustic contamination if, for example, the participant shifted posture. These linear models were fit using the scikit-learn library’s (Pedregosa et al., 2011) *SGDRegressor* class with default parameters save for regularization strength (alpha = 400), early stopping, and initial learning rate (eta0 = 0.0000004). Hyperparameter values were identified based upon manual tuning. Regressions were fit using voltage activity occurring during voicing epochs. These timepoints were identified by extracting the acoustic envelope using a Hilbert transform and then thresholding based upon a manually identified threshold.

### 2.11 Tuning fork control for microphonic pickup

We performed this control experiment during a separate research session with participant T5, twelve months after his primary Many Words Task session. The participant was seated in his chair, facing the microphone, and connected to the full recording system in the same way as during the primary data collection. The participant was asked to sit quietly, relax, and avoid moving while a researcher activated a tuning fork (by striking it with a mallet) and then held it in the air just in front of the participant’s head. The researcher then touched the tuning fork’s stem against T5’s head, then against each Blackrock cable pre-amplifier, and finally against each of the hanging cables itself. The participant reported that he could hear and/or feel the vibrations in all conditions but that they did not cause any discomfort.

We identified 7 second audio and neural snippets corresponding to when the tuning fork was activated and applied to the head/pre-amplifier/cable. Power spectrograms were calculated for these snippets’ raw voltage signals (sampled at 30,000 Hz with no additional filters applied except for the analog filters of the Blackrock Neural Signal Processor) using a short time Fourier transform (50 ms windows, 1 ms sliding window, 7.3 Hz resolution).

## 3. Results

### 3.1 Single electrode modulation when producing speech

We first assessed single electrode phoneme tuning properties by measuring firing rate changes between the instructed delay period (when participants first read a short word) and around speech onset (when they spoke it out loud in response to a go cue). For each electrode, we examined a 500 millisecond window of delay period activity just prior to the go cue and a 100 millisecond window centered on each phoneme’s voice onset time. We then calculated average firing rates within each window (**Fig. 2A**) and tested the significance of FR changes across these two time windows (permutation sign-test, 1000 permutations; uncorrected) for all phoneme classes (**Fig 2B, D**). To maintain statistical independence of samples, if the same phoneme class occurred multiple times within a trial, we randomly selected only one occurence. Nearly all electrodes showed firing rate modulations to speaking at least one phoneme (156/158 electrodes for T5, 179/179 for T11). A sizable minority of electrodes showed tuning to at least half of all phoneme classes (40/158 electrodes for T5, 41/179 for T11, **Fig. 2E-F**). Across electrodes and phonemes, significant firing rate changes largely reflected increasing firing rates around voice onset (78% and 86% of significant changes for T5 and T11 respectively). Thus, a dominant feature of these data was broadly tuned firing rate increase after the go cue, consistent with our previous intracortical multielectrode recordings (Stavisky et al., 2019) and prior ECoG recordings in ventral sensorimotor cortex (Bouchard et al., 2013; Mugler et al., 2018).

**Figure 2.**
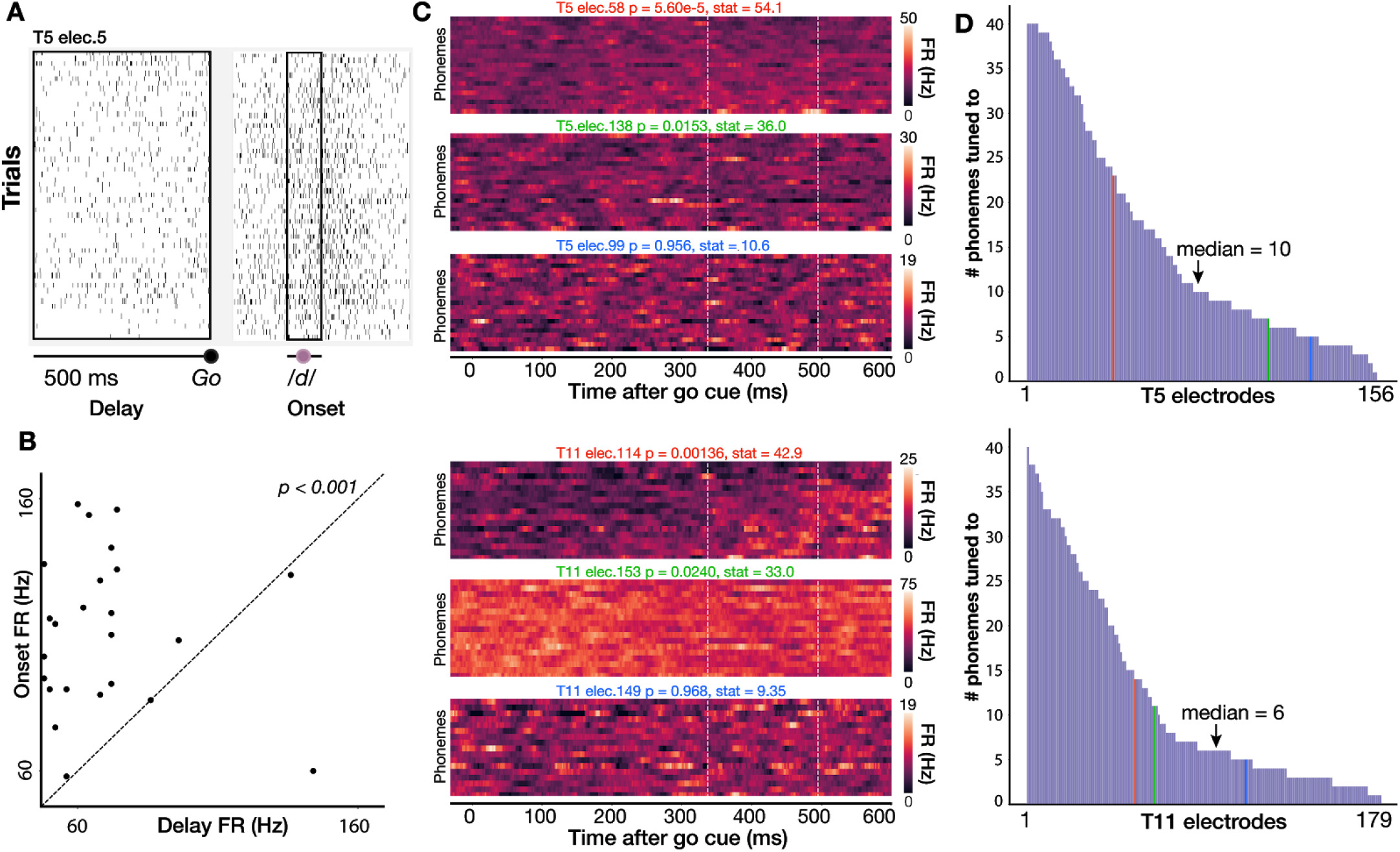
Individual electrodes show broad tuning across phonemes. (**A**) Spike rasters for a single T5 electrode across all instances of /d/ in the full dataset of spoken words. Black boxes show a 500 ms delay period analysis window before the go cue and a 100 ms analysis window centered around voice onset). (**B**) Scatter plot of firing rates during the delay and onset epochs for the electrode shown in A; each point is one trial. Firing rates are significantly higher around voice onset (two-sided permutation sign test; p<0.001). (**C**) Three example T5 electrodes (top) and T11 electrodes (bottom) chosen to exemplify high, low, and not significant selectivity between speaking different phones (Kruskal-Wallis across single trial firing rates from 350 to 500 ms after go cue, marked by vertical lines).. The phonemes were sorted by firing rate for each participant’s high tuning example electrode and then kept in the same order for the other two electrodes. (**D**) Distribution of number of phonemes to which T5’s (top) and T11’s (bottom) electrodes are tuned (i.e., having a significant firing rate difference between delay and onset epochs), sorted from broadest tuning to most narrow tuning. In general, electrodes show a broad tuning profile. Vertical colored lines indicate the corresponding color’s electrode in panel C.

These data did not contain strong preparatory activity or phoneme discriminability during the instructed delay period. Therefore, in the rest of this study we focus on decoding the activity immediately preceding and during overt speaking. Speech epoch firing rates (averaged across electrodes and a 1 s window centered on each phoneme utterance’s onset) were 18.1 ± 35.0 Hz for T5 (mean ± s.d.) and 10.6 ± 27.6 Hz for T11.

### 3.2 Decoding English phonemes using neural population activity

We next sought to assess the utility of these signals for a speech BCI by decoding (offline) the identity of individual phoneme utterances from the neural activity across all functioning electrodes. We trained logistic regression classifiers with neural data and phoneme labels across all utterances (~14 total minutes data) and assessed cross-validated classification accuracy across 39 phonemes. Our first major observation was that although TC-spikes contained substantial information about phoneme identity (14.8% accuracy for T5, 7.9% accuracy for T11, p < 0.002 compared to chance), classification was much more accurate when decoding high-frequency local field potential power (HLFP, 125-5000 Hz): 29.29% and 11.22% for T5 and T11 respectively (**Fig. 3A,F**). This is consistent with our previous study decoding a small number of syllables spoken by T5 (Stavisky et al., 2018) and previous arm and hand BCI studies (Bouton et al., 2016; Pandarinath et al., 2017; Zhang et al., 2018) employing Utah arrays. The superior performance of HLFP decoding may reflect this neural feature more robustly capturing spiking activity from more neurons in the local vicinity of each electrode (Waldert et al., 2013). Since HLFP decoding outperformed TC-spikes decoding, for the remaining analyses we will focus on this more informative neural feature.

**Figure 3.**
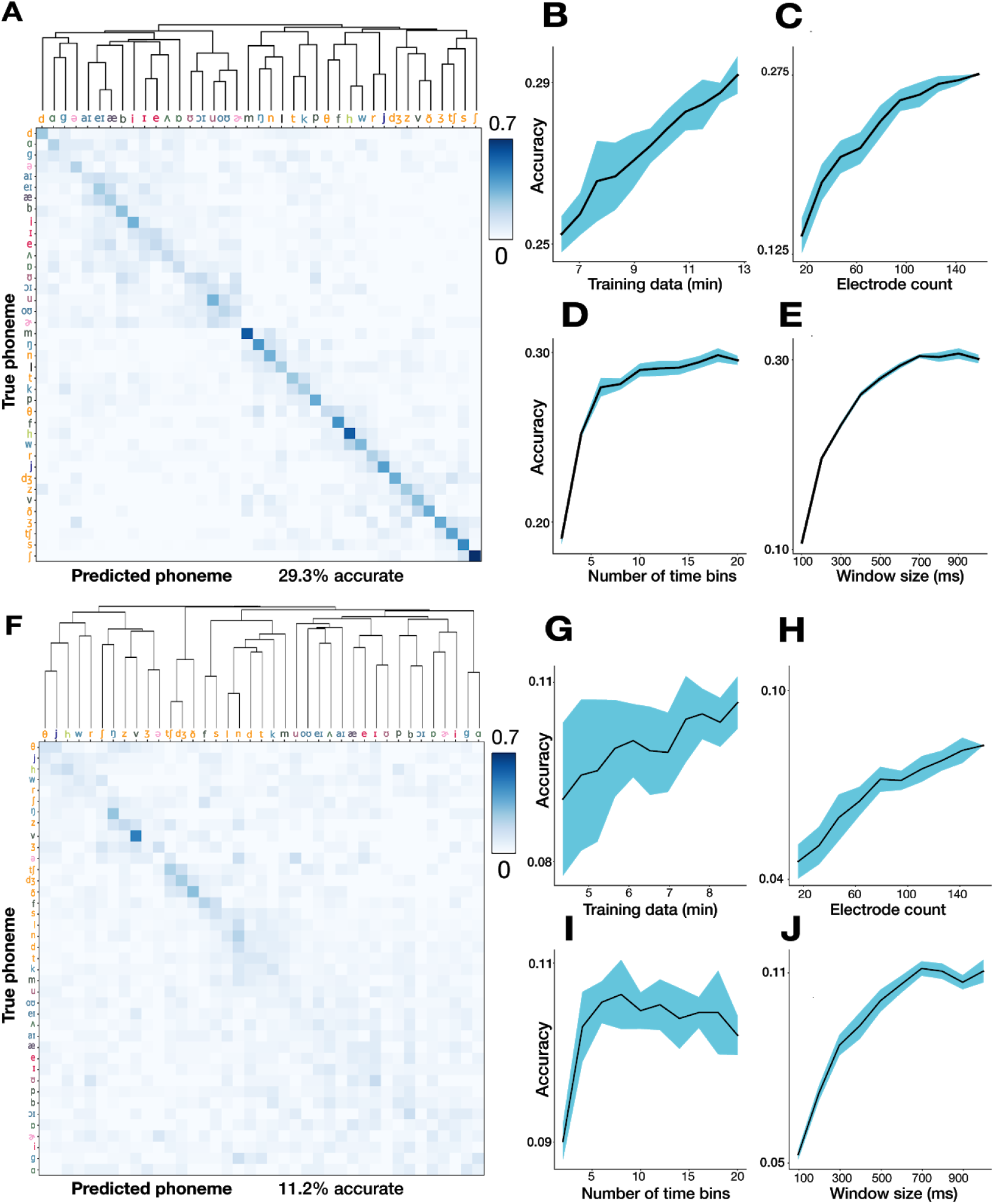
Decoding 39 English phonemes and associated hyperparameter sweeps. (**A**) T5 phoneme decoding confusion matrix, sorted by a hierarchical clustering dendrogram. Values are normalized so that each row adds up to 1. Overall accuracy was 29.3% using leave-one-out cross-validation. Note that the colorbar saturates at 0.7 (to better show the pattern of errors), not 1. Phoneme labels are colored based on their place of articulation group, which is examined further in Supplementary Figure 2. (**B-C**) Parameter sweeps for training set size (B) and number of electrodes (C). Shading denotes standard deviation across 10 repetitions of 10-fold cross-validation. **(D)** Creating finer-grained time bins from the overall 500 millisecond window improves performance. For example. twenty time bins (the rightmost point of this plot) means that each electrode contributes twenty bins each averaging HLFP across 25 ms to the overall neural feature vector. **(E)** Using a larger window (with 50 ms non-overlapping bins) increases performance until saturation around 600 ms. (**F-J**) same as above for T11 data.

To better understand how phoneme decoding varied as a function of the quantity and temporal granularity of the input intracortical data, we measured performance as a function of increasing training dataset size, electrode count, and the number of time bins within an overall 500 millisecond window. Increasing accuracy as a function of both more training data (**Fig. 3B,G**) and more electrodes used (**Fig. 3C,H**) did not show performance saturation. Importantly, this indicates that a path to improve intracortical speech BCI performance would be to collect more data and implant more electrodes. Taking advantage of time-varying structure by dividing each utterance’s neural window into more time bins substantially increased performance (**Fig. 3D,I**). This observation is in accordance with prior ECoG decoding work (Ramsey et al., 2018). Unlike T5’s almost monotonic improvement with more fine-grained time bins, T11’s classification performance saturated at 10 bins (each 100 ms long) and then declined. This trend likely follows from the lower SNR (and lower overall firing rates) in T11’s recordings, in which case short time bins will exacerbate noise in the inputs and reduce decoding accuracy. Using a total time window of approximately 600 milliseconds maximized performance; this agrees with a similar analysis by Mugler and colleagues (Mugler et al., 2014). However, this increasing classification accuracy when using longer time windows should be interpreted with caution for at least two reasons. First, given the limited dictionary size in this study (420 words), increasing the decoding window may allow the classifier to exploit neural correlates of adjacent phonemes in a way that is not representative of what could be expected during open vocabulary decoding. Second, overly long time bins may introduce problematic delays if used in a closed-loop speech BCI (e.g., by delaying when decoded sounds/words are synthesized/typed).

Although phoneme prediction was above chance in both participants, classification accuracy was much higher in T5 than in T11. This result is consistent with ongoing efforts to characterize these T11 signals’ relationship to a variety of actualized and attempted movements (including attempted arm and hand movements), where preliminary findings indicate that T11’s arrays modulate substantially during movements, but exhibit less specificity than the T5 recording. This could reflect differences either in array placement or in modulation specific to this area of cortex in this participant. Given this lack of specificity, we believe that T5’s signals more closely approximate those that would be available in ventral areas of motor cortex that would be recorded from in future work, specifically centered on building a speech prosthesis. Thus, subsequent analyses exploring phoneme decoding in more depth are restricted to the more informative T5 dataset.

To more directly compare the information content in T5’s intracortical recordings to a prior ECoG study reporting what is (to our knowledge) the highest accuracy in decoding phonemes within stand-alone short words (Mugler et al., 2014), we attempted a closely matched comparison to that study. This was facilitated by our having deliberately used a similar set of prompted words to that study: 232 of our words were also used by Mugler and colleagues (Mugler et al., 2014), and we picked replacements for the missing 80 words by choosing similar words from our (larger) word set (see *2.2: Many words task*). For this analysis we also restricted our trials to match the trial count of the participant with the highest performance in Mugler and colleagues (Mugler et al. 2014) and used an equivalent 600 millisecond window centered on voice onset (**Supplementary Fig. 2**). Decoding vowels and consonants separately as in (Mugler et al., 2014), the T5 dataset accuracy was 36.8% across 24 consonant classes (vs. 36.1% in Mugler et al.’s best participant) and 27.7% across 16 vowels classes (vs. 23.9%).

In the previous analyses we used logistic regression decoding in the interest of pragmatic considerations (compute time for parameter sweeps) and facilitating comparison to previous work (Mugler et al., 2014). However, we recognize that additional performance may be gained by employing modern and powerful relatively-new machine learning techniques. To that end, we also decoded T5’s phonemes dataset using a recurrent neural network with data augmentation (see 2.8 *Recurrent neural network decoder*). This resulted in a modest performance improvement (33.4% accuracy) (**Supplementary Figure 3**).

### 3.3 A neural realignment strategy to correct for systematic voice onset biases between phonemes

We next turn to the first of two potential confounds by which studying audible speech may artifactually inflate the accuracy of decoding the corresponding neural data. Recall that, as per standard practice, we defined the start of each phoneme utterance based on the audio recording. This approach makes an important assumption: insofar as detectable speech sound lags the start of the underlying process of speech production, this lag is consistent across different phonemes. However, this assumption is unlikely to hold under most models of what motor cortex encodes. Specifically, if neural activity reflects articulatory movements (Chartier et al., 2018; Lotte et al., 2015; Mugler et al., 2018; Stavisky et al., 2019), then there may be across-phoneme differences between when articulatory movements start versus when the phoneme is clearly audible.

**Figure 4A** presents empirical evidence of this problem by showing different time alignments to generate an example electrode’s firing rates as T5 spoke different phonemes. In the left-most panel, neural data are aligned to the task go cue; note that these analyses only include the first phoneme of each word to facilitate aligning to the go cue or to the neural correlate of speech initiation, as described below. This electrode’s firing rate increased shortly after the go cue and was largely similar across phonemes, consistent with the presence of a large condition-invariant signal, CIS (Kaufman et al., 2016), upon speech initiation (Stavisky et al., 2019). In addition to this general activity increase, there was also phoneme-specific information as indicated by significant firing rate differences between phonemes in the epoch from 350 ms to 500 ms after the go cue. These data also suggest that there were not large systematic differences in the reaction time when speaking words that start with different phonemes. In stark contrast, the **Fig. 4A** center panel shows the same trials aligned to audio-derived voice onset time. The firing rate traces for plosives (/p/, /b/, /k/, /g/, /t/, and /d/) are shown with warm colors to better highlight a systematic onset timing bias. These phonemes’ firing rates follow a similar time-course to the other phonemes’, except for a time offset, which leads to much greater differences across phonemes in an analysis epoch of the same duration (150 ms) now centered on voice onset. These plosive phonemes involve a temporary constriction of airflow (which produces little sound), followed by a rapid (and very audible) release of air, thereby increasing the latency between when phoneme production starts (e.g., movement of the lips) and when the voice onset is detected. This example vividly illustrates that a consequence of using audio data to mark the start of each phoneme utterance introduces systematic timing biases that appear as spuriously accentuated differences across phonemes.

**Figure 4:**
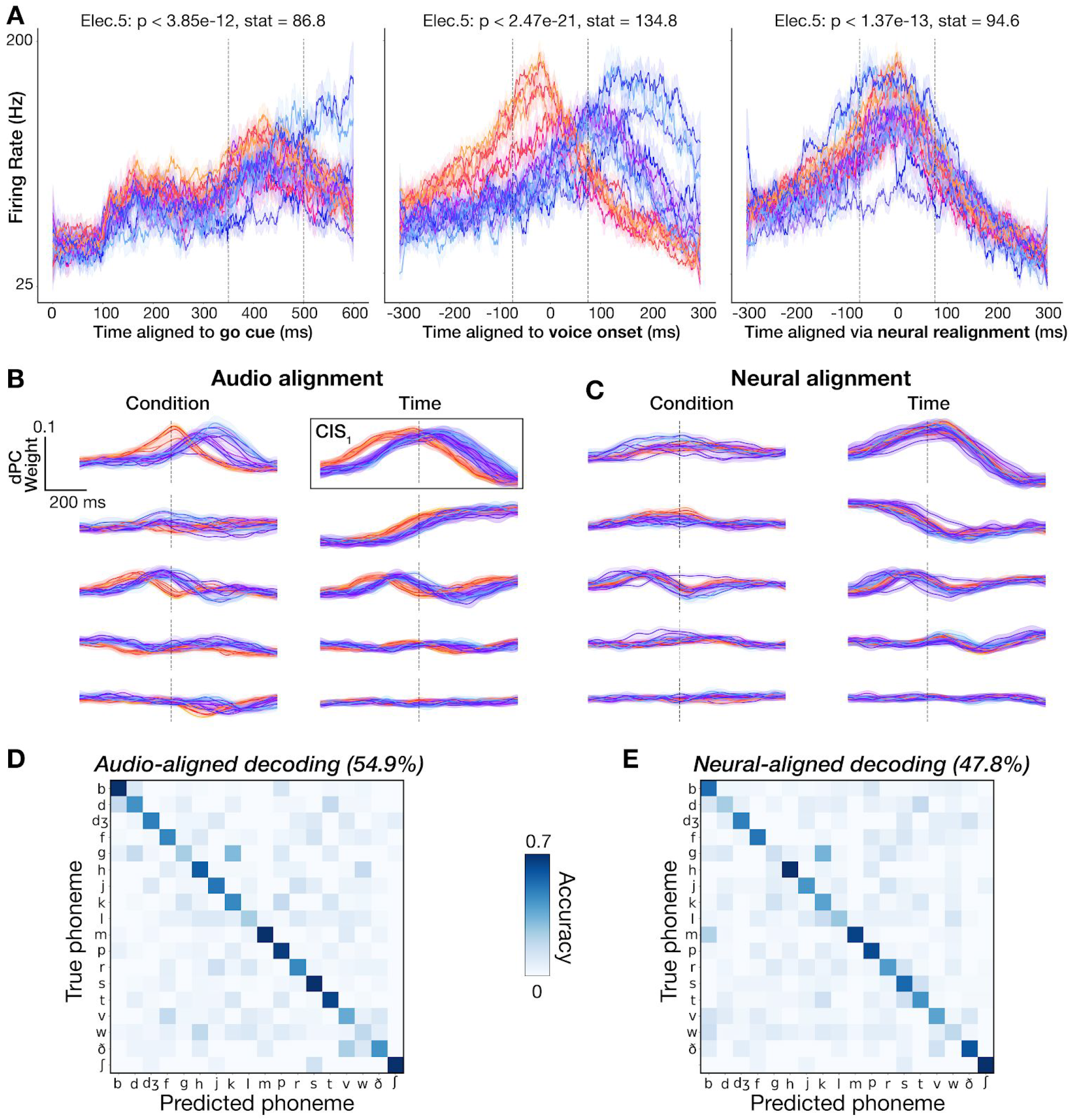
Audio-based phoneme onset alignments cause spurious neural variance across phonemes. (**A**) Firing rate (20 ms bins) of an example electrode across 18 phoneme classes is plotted for distinct alignment strategies (left to right: aligning the same utterances’ data to the go cue, voice onset, and the “neural onset” approach we introduce). Each trace is one phoneme, and shading denotes standard errors. Plosives are shaded with warm colors to illustrate how voice onset alignment systematically biases the alignment of certain phonemes. (**B-C**) dPCs for phoneme-dependent and phoneme-independent factorizations of neural ensemble firing rates in a 1500 ms window. The top five dPC component projections (sorted by variance explained) are displayed for each marginalization for the audio and neural alignment approaches. (**B**) dPC projections aligned to voice onset (vertical dotted lines). Plosives (warm colors) have a similar temporal profile to other phonemes (cool colors) except for a temporal offset. Each phoneme’s trial-averaged peak time of the largest condition-invariant component, outlined in black, was used to determine a “neural onset” for neural realignment. (**C**) Recomputed dPC projections using this CIS_1_-realigned neural data. Vertical dotted lines show estimated CIS_1_ peaks. (**D**) Decoder confusion matrix from predicting the first phoneme in each word using a 500 ms window centered on voice onset. (**E**) Confusion matrix when classifying the same phoneme utterances, but now using neurally realigned data.

As an alternative to audio-derived voice onset, we attempted to detect phoneme onsets directly from the neural data. We used the largest variance phoneme condition-invariant signal of the neural population signal, the ‘CIS_1_’, as a speech activity onset indicator. This linear readout was previously found to be time-locked to attempted motor activity (Kaufman et al., 2016), including during speaking (Stavisky et al., 2019). Since the CIS_1_’s temporal profile is largely invariant across speaking different phonemes, we can perform a “neural realignment” by aligning CIS_1_ peaks across conditions. To do so, we used the demixed PCA (Kobak et al., 2016) dimensionality reduction technique to decompose and summarize the neural ensemble activity into a smaller number of condition-invariant and condition-specific components. **Figure 4B** shows ten of these population modes. The time-courses of these components across phonemes appear very similar except for temporal offsets, providing a population-level corroboration of the previous individual electrode example. Once again, the early peaking traces correspond to plosives.

We then used each phoneme class’ CIS_1_ peak to identify the offset from audio voice onset. This allowed us to realign the corresponding neural data to this ‘neural onset’. For instance, /b/ had an offset of +20 milliseconds; thus, for all trials with words that start with /b/, we shifted our neural data 20 milliseconds forward in time. Performing dPCA again on these realigned data (**Fig. 4C)**, both the condition-invariant and the condition-specific components were more similar across phoneme classes. The right-most plot of **Fig. 4A** shows the same example electrode, now realigned using this CIS_1_-derived neural onset time. Here the time-courses of each phoneme appear more similar than when aligning to audio-derived voice onset, but there were still significant differences between phonemes.

### 3.4 Classification is weakly affected by audio alignment-based phoneme onset timing bias

In the previous section we identified a phoneme timing bias from audio-derived onset labeling, and also demonstrated a method to correct for this bias (at least for the first phoneme of a word) using neurally-derived onset labeling. This allows us to now quantify how much phoneme onset biases can affect decoding audio-labeled phonemes by enabling a classifier to take advantage of spurious neural variance across phoneme classes.

To measure the impact of this confound, we trained separate decoders to classify the first phoneme of T5’s spoken words using 1) the original neural data windows, which were aligned to voice onset as in **Fig. 3**, and 2) the realigned neural data windows obtained using CIS_1_ peak alignment. For both procedures, we used the same preprocessing steps as before (see *3.2*). Decoding each word’s first phoneme from neural data time windows centered on neurally-derived phoneme onsets yielded 47.8% accuracy across 18 classes **(Fig. 4E**), as compared to 54.9% using windows centered on audio-derived phoneme onsets (**Fig. 4D**), a relative decrease of 12.9%. This indicates that that audio-derived phoneme onset biases inflate overt speech decoding accuracies, but only modestly so.

### 3.5 Classification is weakly affected by microphonic artifact and is possible before voice onset

The second potential confounds stemming from studying neural correlates of audible speech is that speaking could mechanically jostle components of the electrophysiology recording system, which would interact with ambient electromagnetic fields to generate small electrical currents that affect the final voltage measurement (i.e., microphonic pickup). This could be a larger or smaller effect, or no effect at all, depending on electromagnetic shielding (e.g., of cables), ambient electromagnetic power (e.g., other powered devices nearby, including lighting) and how recording components are stabilized (e.g., cables immobilized or not). There is also the possibility that there is an acoustic effect, but that it arises from some other mechanism that is not understood. To identify if audio contamination was potentially impacting classification performance, we implemented four different analyses.

We first looked for frequency-specific correlations between audio and neural signals, as in (Roussel et al. 2019). We computed spectrograms for simultaneous speech audio (microphone) recordings and each electrode’s activity (**Fig. 5A**). This yielded a set of time series containing frequency-specific signal power from 5 to 1200 Hz. We isolated time snippets with audible speech (see 2.9: *Quantifying acoustic artifact and Linear Regression Reference (LRR) decontamination)* and correlated these electrode and audio power spectral density (PSD) time series at each frequency . If microphonic pickup had heavily contaminated our recordings, then the most prominent audio frequencies would most likely induce oscillations in the electrode signal at the corresponding frequency and manifest as stronger correlations with the associated “neural” signal power at those frequencies. Plotting the mean PSD across speech timepoints alongside our electrodes’ correlations for an example block, we found strong electrode-audio correlations across a range of frequencies (**Fig. 5B**) with a max correlation *r* = 0.953 across all 192 electrodes × 240 frequencies = 46,080 comparisons, and a median *r* = −0.002. This implies that our neural recordings contain microphonic contamination in at least some frequency bin(s) on some electrode(s). However, inspecting the example electrode which had the strongest overall audio-neural correlation (**Fig. 5A**), we still see little contamination visible to the eye. This contrasts with the striking obvious contamination in (Roussel et al. 2019). The **Fig. 5B** inset summarizes the correlation with audio across all electrodes, and reveals a small set of electrodes with higher audio-neural correlations, all on the same (medial) array .

**Fig. 5:**
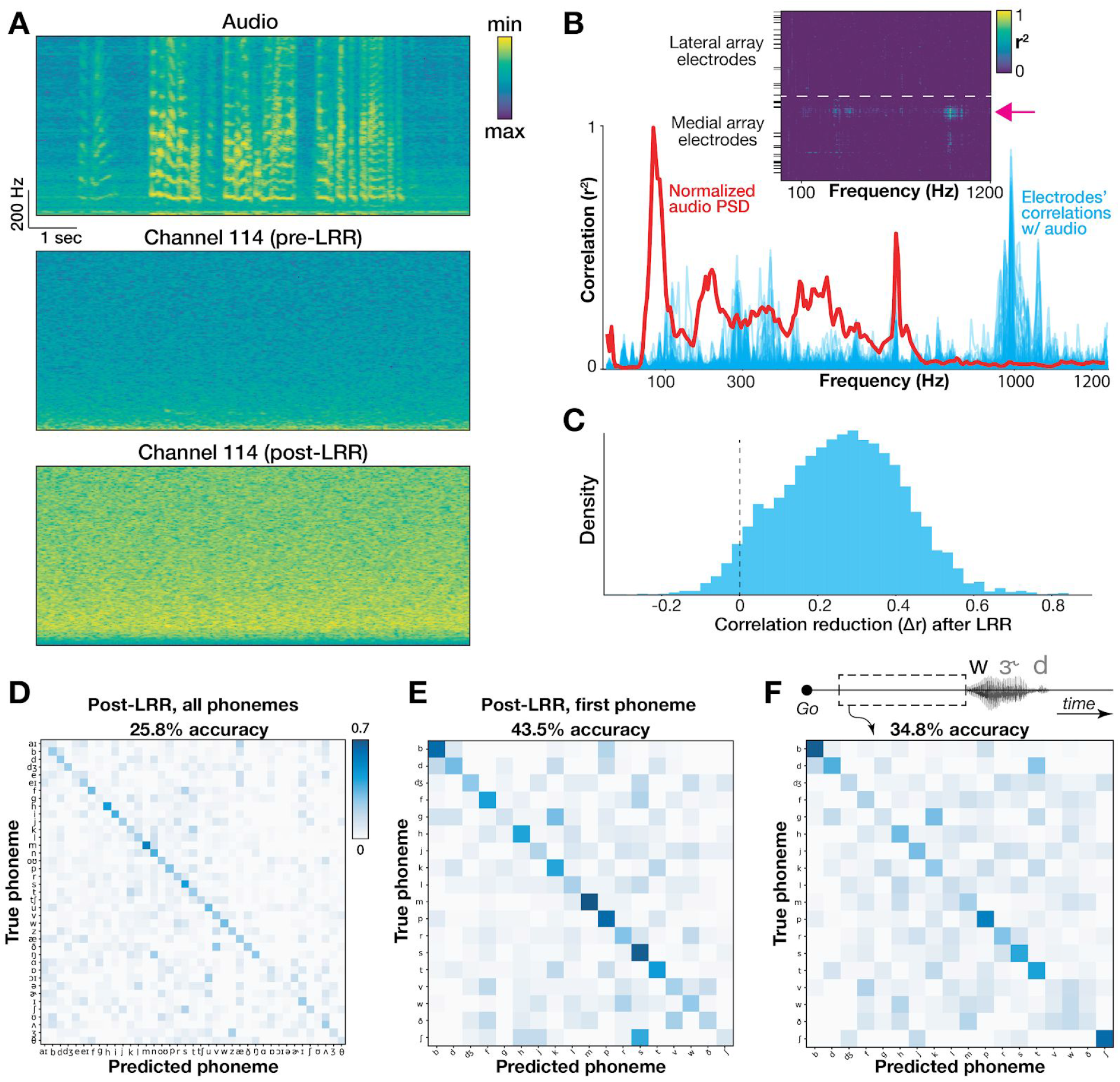
Quantifying and mitigating acoustic contamination of neural signals. (**A**) Spectrograms for audio and neural data in the electrode and block exhibiting the strongest audio-neural correlations. Frequencies range from 5 to 1000 Hz. The bottom plot shows the same electrode after LRR “decontamination”. (**B**) Plot of the mean audio PSD (red) and all electrodes’ Pearson correlations (blue) from the same example block. Inset shows correlation coefficients of individual electrodes (rows) across frequencies (columns). Black horizontal ticks denote electrodes excluded from neural analyses. The pink arrow shows the example electrode from panel A. (**C**) Change in audio-neural correlations after LRR, pooled across all blocks, electrodes, and frequencies (restricted to electrodes with *r*^2^ > 0.1 originally). Values to the right of the dotted ‘0’ line indicate a reduction in correlation strength. The mean audio-neural correlation reduction was 0.26. (**D**) Full classifier confusion matrix after LRR (25.8% overall accuracy across 39 classes). (**E**) Confusion matrix for first phoneme decoding after applying LRR. As in D, the classifier used a 500 ms window centered on voice onset. (**F**) Confusion matrix showing decoding each word’s first phoneme using 500 ms leading up to voice onset to avoid possible audio contamination or neural activity related to auditory feedback.

Second, to more directly test whether mechanical vibrations in the speech range could, in principle, affect our recordings. We performed a positive control where we intentionally applied a mechano-acoustic stimulus to the recording setup. During a separate research session with participant T5, we held an activated 340 Hz or 480 Hz tuning fork either in the air in front of the (fully connectorized) participant (**Supplementary Fig. 4A**), or pressed the activated fork’s stem to his head near the pedestals (**Supplementary Fig. 4B**), directly to the pre-amplifiers (**Supplementary Fig. 4C**), or directly to the connector cables (**Supplementary Fig. 4D**). We reasoned that since a tuning fork applies the majority of its mechanical energy into the recording apparatus at a specific frequency (unlike speech), and is unlikely to evoke biological neural oscillations in a motor cortical area at that same specific frequency, then a sharp increase in recorded signal power at the same frequency would clearly indicate the presence of an artifact. We did indeed observe such an artifact on some medial array electrodes when the tuning fork was applied to the pre-amplifier and cable, the two conditions in which there was putatively the most mechanical energy being transferred into the recording apparatus. This is an important built-in positive control, demonstrating that our measurement and analysis methods do have the sensitivity to detect, at least this level power at this frequency, of contamination. We do not purport to equate this artificial tuning fork stimulation to the acousto-mechanical interaction between the participant speaking and the recording system; rather, this result, which is consistent with (Roussel et al., 2019), merely shows that there exist conditions under which artifictual acoustic contamination is possible. This lends support to the aforementioned suspicion that some of the speech-correlated neural activity was due to acoustic contamination.

Third, as a conservative decoding analysis that would avoid acoustic contamination, we extracted a 500 millisecond neural window just prior to each word but before speech onset. Since this neural activity precedes audible audio signals, it cannot be polluted by microphonic pickup. This analysis also serves as a control that excludes the potential contribution to decoding by neural processing of auditory feedback in this motor cortical area. Decoding these pre-voice neural windows for each word’s first phoneme yielded a cross-validated (20 fold) classification performance of 34.8% across 18 classes (**Fig. 5F**), well above chance performance (p < 0.002; permutation test, 500 permutations). For comparison, using a 500 ms neural window centered on the phoneme’s voice onset (which we expect to better capture phoneme production-specific neural activity, in addition to potential acoustic contamination and auditory feedback) yielded an accuracy of 54.9% (**Fig. 4D**).

Fourth, we implemented a denoising procedure (linear regression reference or ‘LRR’, Young et al., 2018) which aims to mitigate acoustic contamination of the neural signals. LRR works by subtracting out predictable signals shared across many electrodes (which is what we would expect from microphonic pickup) from each electrode’s activity. A linear regression model estimates each electrode’s activity using instantaneous activity from all other electrodes; this prediction is then subtracted from that electrode’s original time series to yield the denoised estimate. This approach was first applied in (Young et al., 2018) to remove electrical artifacts induced during intracortical microstimulation. Here we applied the method to putative acoustic contamination by fitting regression models to bandpass-filtered (125-5000 Hz to match our HLFP neural feature) voltage signals. **Figure 5A** shows an example electrode’s activity before and after LRR. Following LRR, we observed a reduction in audio-neural correlations (**Fig. 5C**) and only a small reduction in cross-validated accuracy decoding all phonemes (reduction from 29.3% to 25.8%, **Fig 5D**). We note that the LRR technique does not guarantee that all acoustic contamination is eliminated (which would cause us to under-estimate the contribution of microphonic pickup to decoding accuracy), and it also can subtract out genuine neural signal that is shared across electrodes (which would cause us to over-estimate the contribution). Thus, these results should be viewed as just one piece of evidence towards estimating the effects of acoustic contamination in overt speech decoding studies.

In summary, our results support the concern raised by (Roussel et al. 2019) that microphonic artifacts are present in intracortical and ECoG recordings during overt speech. These audio-driven signals may well have somewhat increased the phoneme classification accuracies in this study. However, our further analyses described in this section bracket this potential artifactual contribution as small and, critically, show that we can decode phonemes even before audible voicing (**Fig. 5F**). This agrees with our prior findings that neural activity in this area modulates in response to unvoiced orofacial movements (Stavisky et al., 2019).

### 3.6 Classifier confusions correlate with place of articulation groupings

Lastly, we return to the observation that when sorting our decoder confusion matrices based on the results of agglomerative hierarchical clustering (Sokal & Michener 1958), the resulting dendrograms suggest divisions across phonemes relating to their phonemic groupings (**Fig. 3A**; e.g., broad vowel-consonant divisions). A permutation test (see 2.7: *Logistic regression for phoneme classification*) revealed significantly higher confusion within place of articulation groups than between groups (**Supplementary Fig. 5A-B**; p < 0.005), indicating that there was latent structure in our classifier errors putatively related to how similar the motoric demands of producing different phonemes were (Lotte et al. 2015; Mugler et al., 2018; Chartier et al. 2018; Stavisky et al. 2019). However, we identified in sections 3.3 and 3.4 that these classifier errors were somewhat affected by systematic across-phoneme differences in labeled voice onsets. This raises the concern that the pattern of decoding errors could be related to similarities in phonemes’ latencies between speech production initiation and voice onset, rather than place of articulation *per se*. We therefore repeated the place of articulation grouping analysis, but now applied to classifier errors from predicting the first phoneme of each word using neural alignment (as described for **Fig. 4**). We again observed higher confusion within place of articulation groups (**Supplementary Fig. 5C-D**; p < 0.005), suggesting that this error structure was not just an artifact of voice onset bias.

## 4. Discussion

The present results indicate that decoding speech from intracortical arrays is a promising future direction for developing BCIs to restore lost speech. The phoneme classification accuracy demonstrated in the participant with more informative signals (T5) is similar to work performed with ECoG arrays that spanned a wide swath of speech-related cortical areas (Livezey et al., 2019; Mugler et al., 2014). This is despite the limited spatial coverage of the present study’s Utah arrays (10 × 10 grid of electrodes spanning 4.0 mm × 4.0 mm; array backplane is 4.2 mm × 4.2 mm) and their suboptimal placement outside traditional speech areas. This accuracy also exceeds a prior intracortical phoneme decoding study which used fewer electrodes and reported 21% accuracy across 38 phonemes, (Brumberg et al., 2011). However, the lower accuracy in the second participant (T11), and the overall gap between either participant’s decoder performance and the close to 100% accuracy one would aspire to for a speech BCI that approaches a healthy speaker’s capabilities, motivates future work using more electrodes placed in ventral speech areas.

As in prior ECoG studies (Moses et al., 2019; Mugler et al., 2014), we found that classifier errors exhibited articulatory structure (**Supplementary Fig. 5**). Phonemes spoken with similar place of articulation were more likely to be confused with one another compared to phonemes with different places of articulation. This suggests that the underlying neural signals relate to the movement of articulator muscles, broadly in line with ECoG studies (Chartier et al., 2018; Lotte et al., 2015; Moses et al., 2019; Mugler et al., 2014, 2018) and earlier work looking at spiking activity in this dorsal ‘hand knob’ area (Stavisky et al., 2019). Assuming that our signals reflect articulator movements underlying speech, then the observed classification accuracy improvement from using multiple time bins likely reflects the increased ability to discriminate the time-varying movements underlying speech production (Chartier et al., 2018; Mugler et al., 2014; Ramsey et al., 2018). Future decoding approaches that seek to specifically assess and optimize performance with an eye toward eventual speech BCI applications may benefit from classifier cost functions and performance metrics tailored to speech perception, rather than treating all phoneme mistakes the same. For example, if the final BCI output is sound synthesis, then metrics based upon distortions of auditory spectral features (Anumanchipalli et al., 2019) may be more informative: misclassifying similar sounding phonemes (e.g. /p/ and /b/) would be less damaging to overall speech comprehension compared to mistakes across groupings (e.g. /p/ and /r/).

We observed better performance classifying consonants versus vowels (**Supplementary Fig. 2**), which was also seen in earlier ECoG studies (Brumberg et al., 2011; Livezey et al., 2019; Mugler et al., 2014; Ramsey et al., 2018), although the factors explaining this phenomenon may differ for different recording modalities. In our case, lower vowel decoding accuracy might arise from the dataset statistics: most of the words our participants spoke were three phonemes long and consisted of consonant-vowel-consonant, resulting in more consonant training data. A second possibility is that the 500 millisecond decoding window used is short compared to the audio durations of vowels (**Supplementary Fig. 1C**, 319 ms on average for T5); the resulting neural features may therefore cover more of the relevant information for short duration consonants (173 ms on average for T5). Future work might consider a hierarchical classifier that first tries to decode vowel versus consonant, and then uses different window sizes accordingly. A third possibility is that the underlying articulator movements involved in vowel production are less well sampled by these recordings (Brumberg et al., 2011). Definitively testing the latter hypothesis will require future recordings of simultaneous neural and speech articulator activity.

The improved decoding performance when using high-frequency LFP power (as opposed to threshold crossing spikes) suggests that local voltage fluctuations can support phoneme classification without a need for spike detection. Indeed, prior decoding work across a more limited set of syllables found no accuracy difference between HLFP or spike features (Stavisky et al., 2018). While this suggests that broadband LFP features are sufficient for high-performance decoding, we caution that microelectrode field potentials, which are largely dominated by local action potentials (Nason et al., 2020), reflect different underlying physiological sources than surface electrode recordings. Importantly, performance here was dependent on sampling a sufficiently high number of electrodes from the two arrays (**Fig. 2C**). Corroborating this finding, individual electrodes had broad tuning profiles across phonemes, with 25% (T5) and 23% (T11) of all electrodes showing significant tuning to at least half of the 39 phonemes.

Leveraging more recent machine learning approaches, we obtained a modest performance improvement using an RNN (33% accuracy in a 10-fold CV, compared to 29.3% using logistic regression). Our results indicate that performance had not saturated when increasing the number of electrodes or training data, and even more data should further improve classification accuracy, especially for deep learning methods (Livezey et al., 2019). However, it seems unlikely that arrays in the hand knob area would support a very high performance speech BCI, even with more electrodes or training data. Future efforts will benefit from access to ventral speech cortex as well as higher electrode densities that record from a larger neural population.

In this study we also characterized and addressed two confounds that may be endemic to decoding overt speech. The first has not, to the best of our knowledge, been specifically examined before: labeling speech start times based on voice onset can artificially boost decoding performance by introducing systematic timing differences across phoneme classes between the onset labels and the true time when neural activity rapidly changes during speech production. This timing bias can be seen at the level of individual electrodes (**Fig. 4A)** and at the neural population level (**Fig. 4B)**, and is consistent with the neural activity largely reflecting underlying articulatory movements (Bouchard et al., 2013; Chartier et al., 2018; Lotte et al., 2015; Mugler et al., 2018), rather than when these movements cause detectable sounds. A systematic discrepancy between voice and neural onset times was noted in a previous study (Jiang et al., 2016), which also proposed decoders that exploit relative neural onset differences between phoneme-specific and phoneme-independent neural components. It has also been previously noted (Mugler et al., 2014) that phoneme decoding accuracy is sensitive to the precision of onset timing labeling. Here, we realigned phonemes’ analysis windows to an onset time derived from the neural population signals themselves to account for this experimental confound and demonstrated that high performance classification was still possible (47.8% accuracy across 18 classes, **Fig. 4E**). We anticipate that this neural realignment strategy will also work in other recordings that contain strong condition-invariant responses time-locked to the speaking task. Because we observed an appreciable difference between decoder performance with and without this neural realignment (a 12.9% reduction in accuracy when classifying the first phoneme of each word), we caution that future speech BCI work based on voice onset labels should explicitly take across-labels timing biases into account.

We additionally found evidence of a second confound: acoustic contamination in our electrode signals (Roussel et al., 2019). Strong correlations between audio and “neural” activity occurred at specific frequencies, particularly in a cluster of electrodes on our medial array. However, unlike the characteristic correlation peaks around the fundamental frequency (and harmonics) of participants in (Roussel et al., 2019), our correlations exhibited a less consistent relationship with T5’s audio PSD. This discrepancy may arise from the relatively short snippets of speech we had access to (as opposed to continuous speech), or they may reflect differences in the mechanics of acoustic contamination between these studies’ setups. Further supporting our findings, we observed strong vibrations in the relevant frequency bands of some electrodes when applying a tuning fork to both our participant’s pre-amplifier and cable. To account for this contamination, we implemented two different approaches: 1) classification of each word’s first phoneme using pre-voice onset activity, and 2) the novel use of a decontamination procedure previously deployed for electrical stimulation artifact reduction (Young et al., 2018). By decoding phonemes using only pre-voicing neural activity (**Fig. 5D**, 34.8% accuracy across 18 classes), we showed that our performance is not solely a product of acoustic contamination. By mitigating correlations through a re-referencing approach, we found that we could still decode phonemes with relatively high accuracy (25.8% across 39 classes, although see *4.1*). Thus, our conclusions are similar to that of (Roussel et al., 2019): microphonic pickup is a (manageable) nuisance for overt speech decoding studies, but the majority of the measured voltage signals are biological and not artifactual.

### 4.1 Limitations

In this study we classified neural activity snippets aligned to phoneme onset, as determined from the audio recording of spoken speech. A clinical speech BCI will need to identify speech elements (such as phonemes) from unlabeled, continuous speaking data. Recent work decoding continuous speech from ECoG recordings indicates that this is feasible (Anumanchipalli et al., 2019; Herff et al., 2015), and we anticipate that similar methods can be applied to intracortical recordings. While this presents a greater challenge, on the other hand, natural language processing methods may provide substantial help in the continuous regime (Li and Negoita, 2018). In particular, leveraging the statistical structure of language may enable “denoising” of a neural decoder’s initial outputs (Moses et al., 2019; Willett et al., 2020a). No such corrections were employed in this study, where we instead sought to understand ‘raw neural performance’ before applying any language model (the effectiveness of which will depend on the specifics of the speech BCI application, e.g., open vocabulary versus a more constrained vocabulary).

Our neural realignment procedure allowed us to disentangle the contribution of true (neural) variance across phonemes from that of spurious variance driven by systematic onset timing biases. While the results indicate relatively unimpaired performance following realignment (47.8% post-alignment vs 54.9%, 18 distinct classes), this check was restricted to classifying the first phoneme due to unreliable neural timing signals for subsequent phonemes. It thus remains an open challenge to quantify and mitigate the effect of voice onset biases for the later phonemes of a word.

Our recordings show evidence of acoustic contamination, which most likely improves classification performance. While our LRR procedure was able to mitigate this contamination, some acoustic contamination was probably still present. Future directions could examine a related artifact mitigation method (O’Shea and Shenoy, 2018), with which we anticipate similar results. The present correlation analysis also assumes that the process of sound wave conversion into voltage signal fluctuations preserves frequencies, as opposed to shifting them around. While our tuning fork experiment suggests that this is approximately true (**Supplementary Fig. 5** - although this is not necessarily the same mechanism underlying speech-induced acoustic contamination), we note that Roussel and colleagues observed a slight shift in correlation peaks for a participant with intracortical arrays relative to their voicing fundamental frequency.

Additionally, it is possible that neural signals relating to auditory feedback, as opposed to speech articulation, may drive some of the classification performance in this study. However, we were able to decode words’ first phonemes without an audible voicing activity and, in prior work (Stavisky et al., 2019), we also found strong tuning to unvoiced orofacial movements. Together, these findings suggest minimal feedback-driven signals compared to articulatory information. Future studies are needed to definitively disentangle these two processes two during attempted speech decoding.

These confounds complicate direct comparisons to previous work, as their extents may vary between different studies’ recording setups. Given the different corrections that can be applied (auditory feedback restriction, microphonic pickup controls, neural realignment, etc.) and combinations thereof, we therefore caution that the phoneme decoding comparisons made here should not be viewed as exact, matched performance tests but rather ballpark estimates. With these points in mind, the demonstrated performance is at least comparable to prior studies (Brumberg et al., 2011; Livezey et al., 2019; Mugler et al., 2014) and supports further intracortical speech decoding work.

## 5. Conclusions

Taken together, our results indicate that the limited spatial coverage of current intracortical electrode arrays is more than offset by the high speech-related information provided by intracortical recordings. Our offline decode results suggest a lower bound on intracortically-driven speech BCI performance, since these arrays in a “hand/arm” area of precentral gyrus were likely suboptimally placed for speech decoding, and the participants did not receive online feedback that they could use to beneficially adjust their neural activity. This study de-risks and motivates future work in which arrays are implanted into ventral speech cortex in participants who cannot speak.

## Acknowledgements

We thank participants T5, T11 and their caregivers for their generously volunteered time and effort as part of our BrainGate2 pilot clinical trial. We also thank Professor Mark Slutsky for providing the many words list; our Stanford NPTL and NPSL group for helpful discussions; Beverly Davis, Erika Siauciunas, and Nancy Lam for administrative support; and Dr. Darrel Deo (Stanford, NPTL group) for providing RNN decoder code documentation.

Some of the computing for this project was performed on the Stanford High Performance Computing Cluster (Sherlock cluster). We thank Stanford University and the Stanford Research Computing Center for providing expert IT support for this computational resource where our primary compute node is housed.

This work was supported by an NSF Graduate Research Fellowship DGE-1656518 and Regina Casper Stanford Graduate Fellowship (G.H.W.); the A. P. Giannini Foundation, Wu Tsai Neurosciences Institute Interdisciplinary Scholars Fellowship, and Burroughs Wellcome Fund Career Award at the Scientific Interface (S.D.S.); Howard Hughes Medical Institute (F.R.W., D.T.A); US NIH NIDCD R01DC014034, NINDS R01NS066311, NIDCD R01DC009899, NICHDNCMRR N01HD53403, N01HD10018, Department of Veterans Affairs Rehabilitation Research and Development Service B6453R, MGH Deane Institute for Integrated Research on Atrial Fibrillation and Stroke (L.R.H.); NIDCD R01-DC014034, NIDCD U01-DC017844, NINDS UH2-NS095548, NINDS UO1-NS098968, Larry and Pamela Garlick, Samuel and Betsy Reeves, Wu Tsai Neurosciences Institute at Stanford (J.M.H and K.V.S); NIBIB R01-EB028171 (S.D.); Simons Foundation Collaboration on the Global Brain 543045 and Howard Hughes Medical Institute Investigator (K.V.S).

---

**Supplementary Figure 1.**
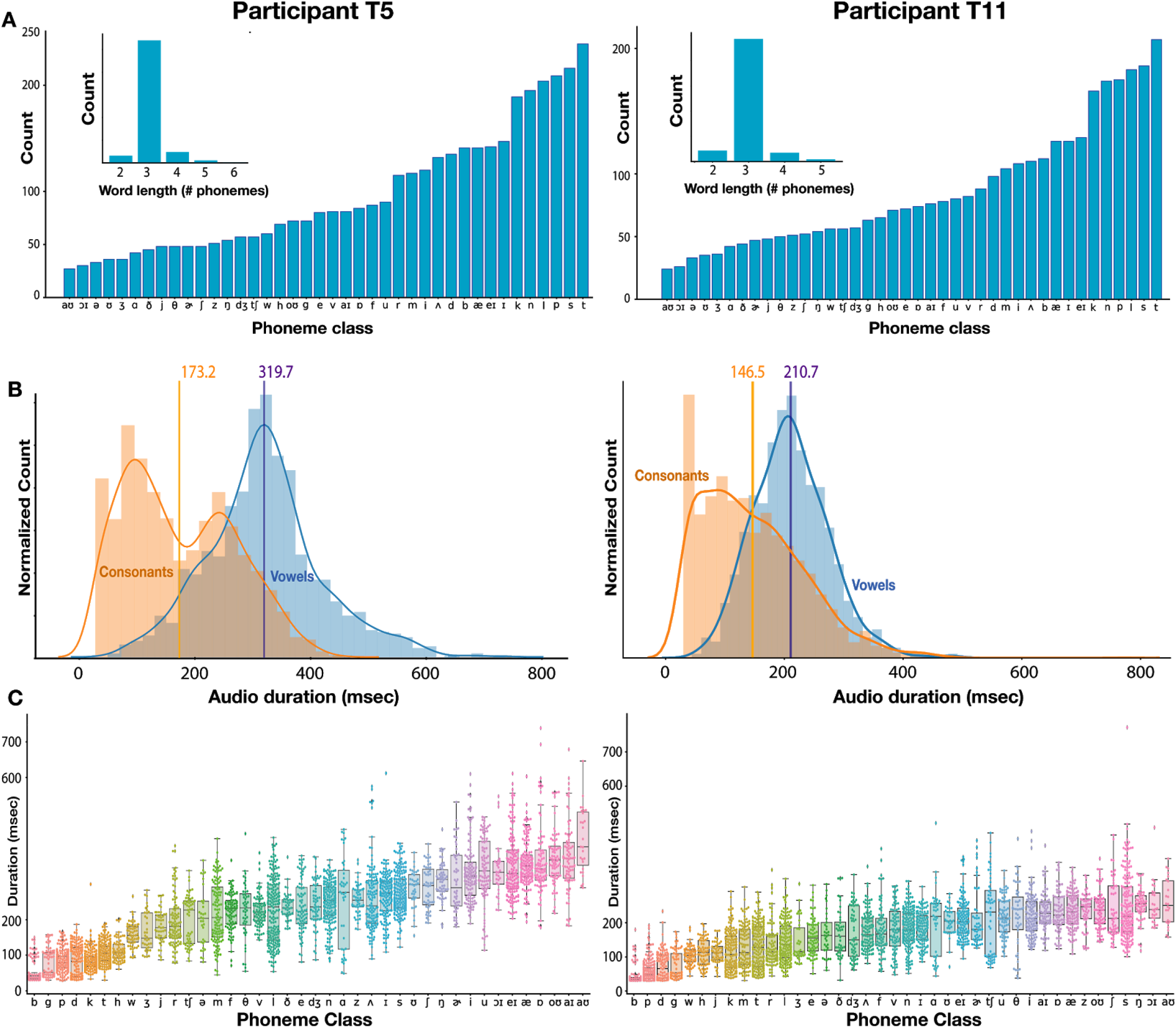
Spoken word and phoneme statistics. **(A)** Distribution of different phoneme class frequencies (T5 - min: 27, max: 239; T11 - min: 24, max: 207). The exact utterance distributions differ between participants due to occasional missed trials or misspeaking. Insets show the distribution of word lengths. A majority (87% for T5, 85% for T11) of words are 3 phonemes long. **(B)** Distribution of phoneme audio durations in milliseconds, split by vowels and consonants. Vertical lines with number labels show each class’ mean. Vowels are longer on average. **(C)** Distribution of phoneme durations, broken down by specific class. Box plots display the median (middle horizontal lines), interquartile range (upper and lower lines), and outliers (Lilliputian diamonds).

**Supplementary Figure 2.**
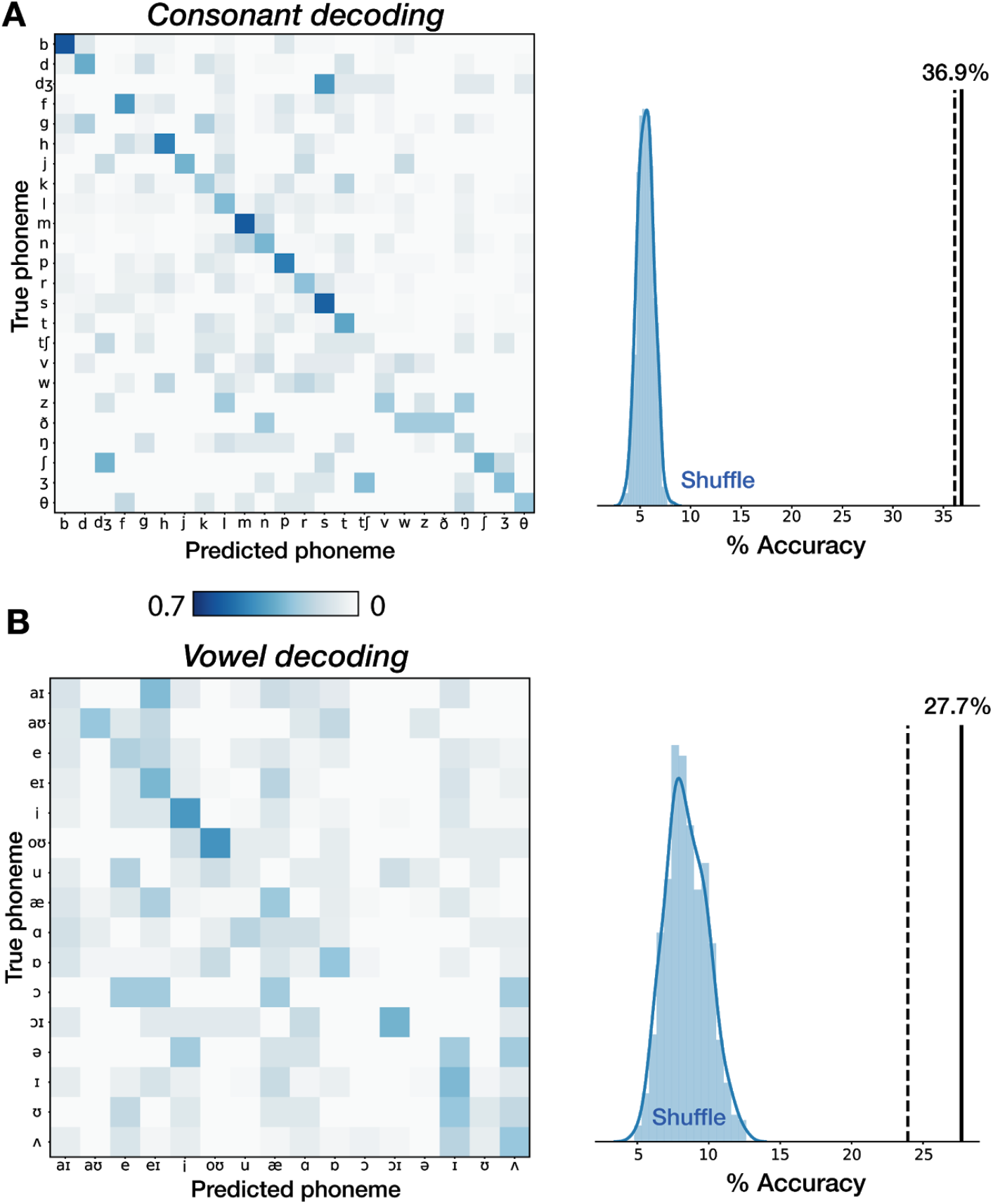
Comparing decoding accuracy to a previous ECoG study with similar spoken words Phonemes were classified using a 600 ms neural window with 50 ms, non-overlapping bins. Cross-validated decoding accuracy across 20 folds is reported. (**A**) Confusion matrix when decoding 24 consonants using a phoneme set closely matched to (Mugler *et al.* 2014). Measured intracortical decoding performance (36.8%, solid line) is comparable with that of Mugler *et al.* 2014 (36.1%, dashed line) and well above chance (p < 0.002; permutation test, 500 permutations). (**B**) same as (A) but with 17 vowels. Measured performance (27.7%) is comparable with that of Mugler *et al.* 2014 (23.9%) and well above chance (p < 0.002; permutation test, 500 permutations).

**Supplementary Figure 3.**
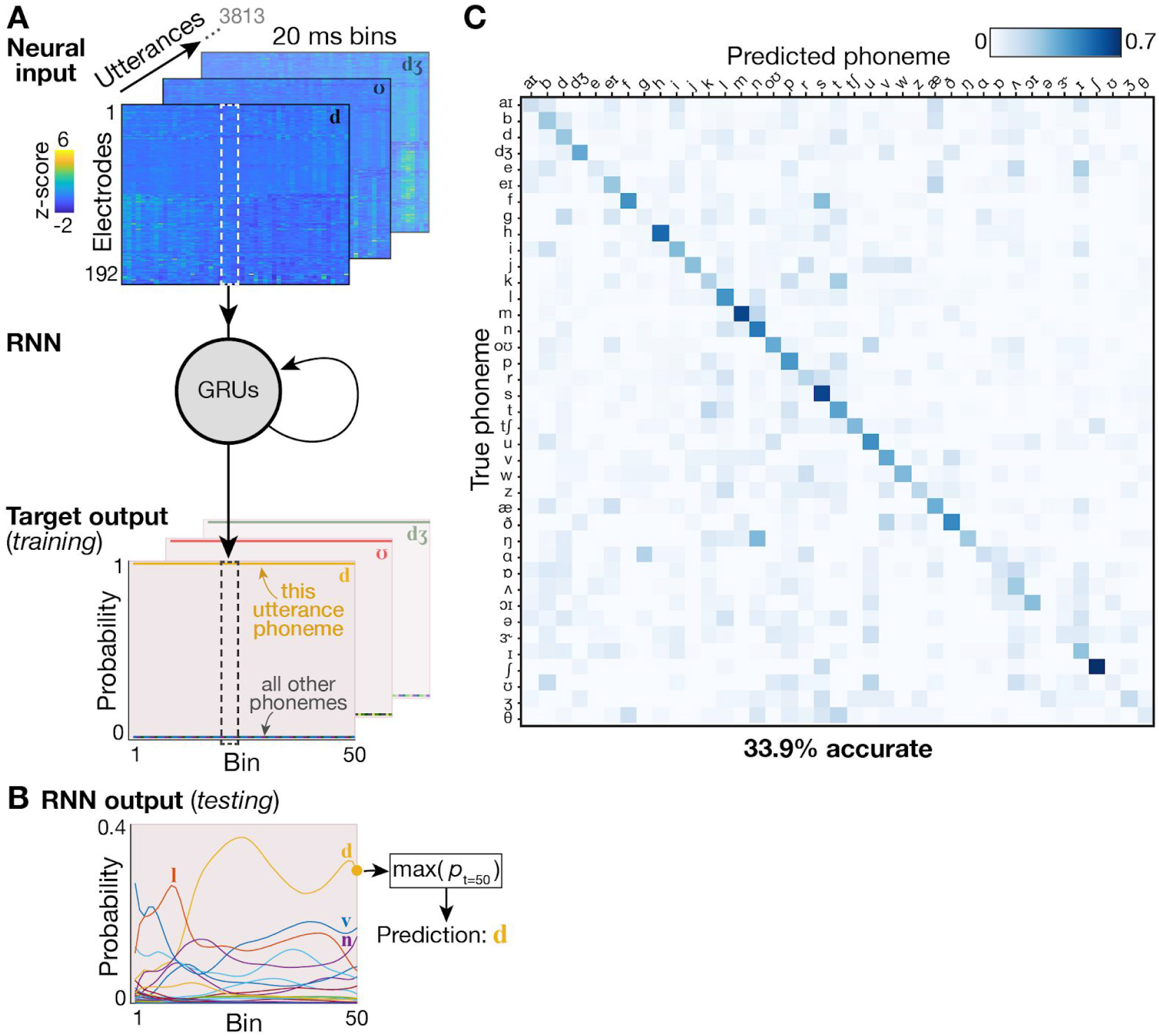
Phoneme decoding using a recurrent neural network (RNN). **(A)** Schematic overview of the RNN decoding approach. A single-layer RNN consisting of 512 gated recurrent units (GRUs) was trained to map neural inputs (top) at each time step to a ones-hot output (bottom) representing which phoneme utterance this neural data snippet came from. Each utterance provided one input snippet consisting of 1000 ms of HLFP activity per electrode (divided into fifty 20 ms bins) centered on the phoneme onset. **(B)** Example network output when the RNN was provided held-out neural data as input. Although estimated probabilities for each phoneme are read out during every time bin, for the utterance’s final discrete output we used the most probable phoneme at the last time bin (here, that would be the phoneme /d/). **(C)** Cross-validated confusion matrix using the RNN to classify all of T5’s phonemes. The 33.9% overall accuracy is slightly improved compared to the 29.3% accuracy when using a linear decoder (see Fig. 3).

**Supplementary Figure 4.**
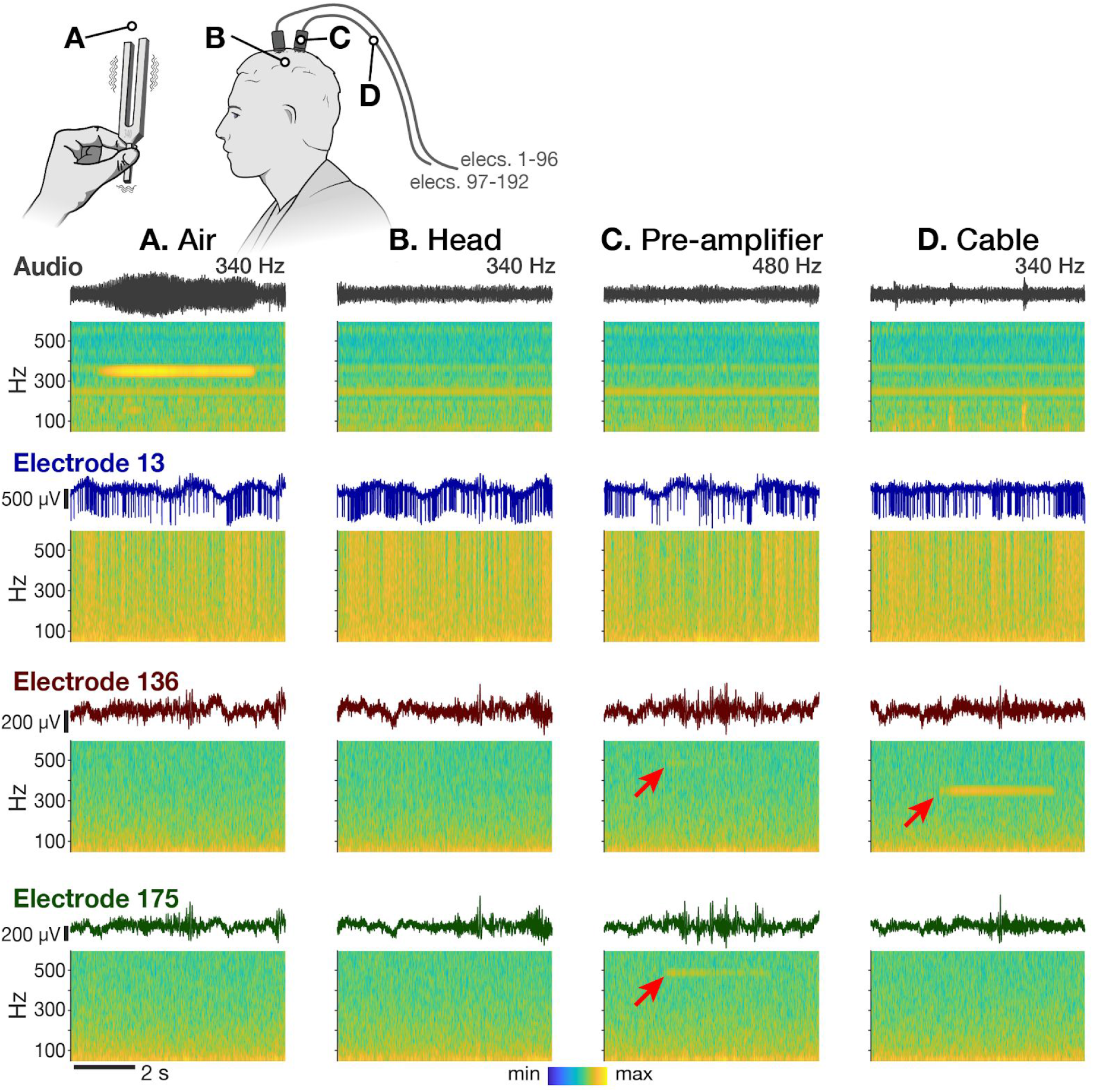
Neural, tuning fork recordings reveal microphonic pickup. In this positive control, a 340 Hz or 480 Hz vibrating tuning fork was held in the air (**A**), gently pressed against participant T5’s head (**B**), pressed against the pre-amplifier (**C**), or pressed against the cable (**D**). The top row shows a snippet of audio channel recording and corresponding acoustic power spectrogram for each condition. The remaining rows show simultaneous raw voltages and and power spectrograms from three example electrodes. Electrode 13 was chosen as an example with no apparent microphonic artifact (it also has prominent action potentials). Electrode 136 showed a microphonic artifact at the tuning fork’s frequency (marked with a red arrow) in the pre-amplifier (C) and cable (D) vibration conditions. Electrode 175 showed an artifact in the pre-amplifier (C) condition, but not the cable condition. We only observed this artifact on the medial array (electrode numbers ≥ 97). The artifact could be generated with either the 340 Hz or 480 Hz tuning forks applied to either of the two pre-amplifiers/cables, but was stronger when applied to the medial pre-amplifier and cable, as in the examples shown.

**Supplementary Figure 5.**
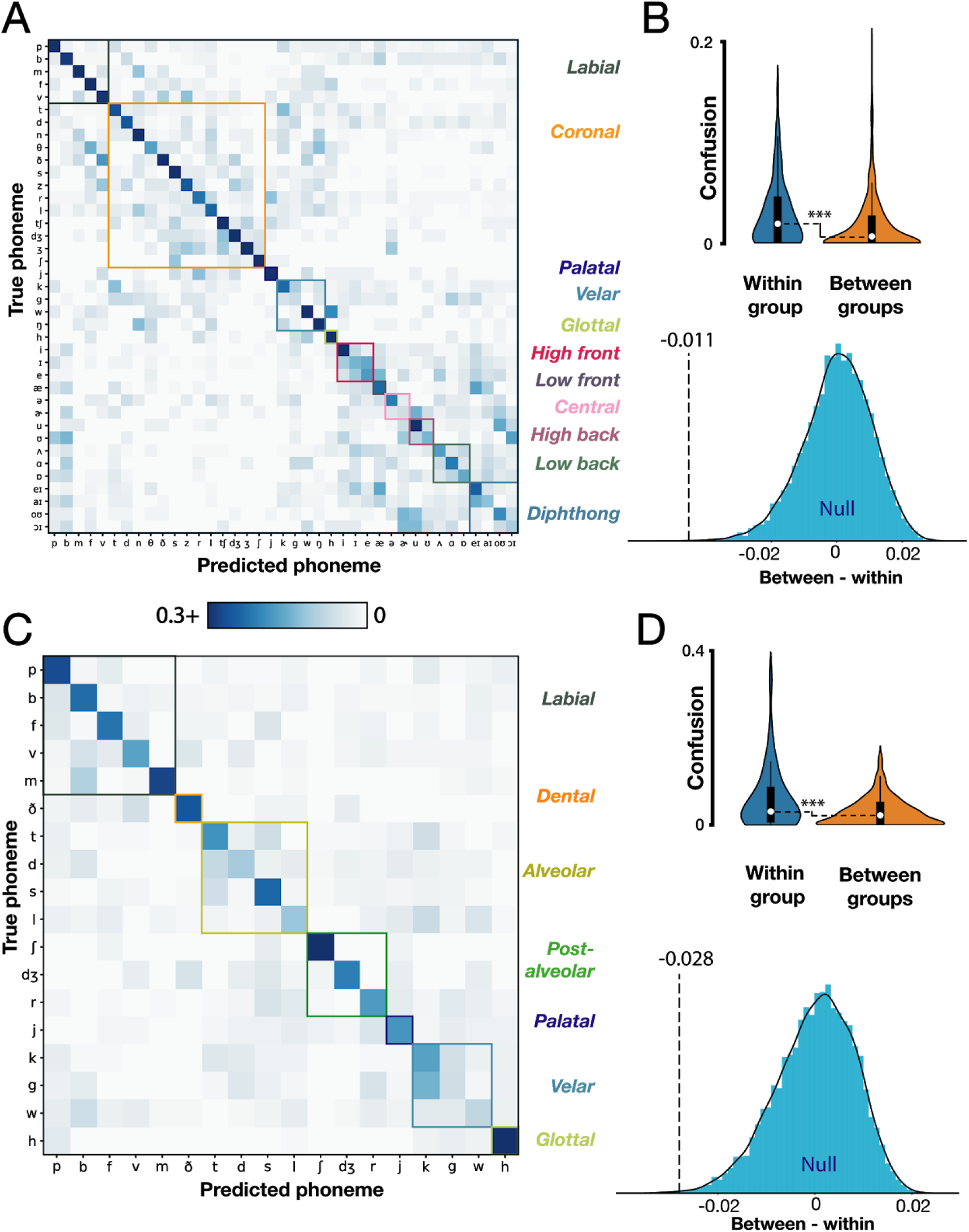
T5 classifier errors reflect articulatory groupings of phonemes. **(A)** Decoder confusion matrix sorted by articulatory groups. Note that the color axis is rescaled to [0 : 0.3] to highlight confusion patterns. **(B)** top: empirical within- and between-group normalized errors (Gaussian kernel densities, box plots with medians and interquartile ranges overlaid). Comparison of within and between group means reveals significantly higher confusions within groups (p < 0.0005); bottom: corresponding null distribution when articulatory groupings are shuffled (blue) compared to the true empirical difference (black dotted line). **(C)** Confusion matrix for predicting neurally-realigned first phonemes of each word. **(D)** Associated significance testing results, presented as in panel B. A permutation test revealed higher confusion within groups (p < 0.002) even after correcting for the biases in audio-derived voice onsets.

